# SARS-CoV-2 RNA reverse-transcribed and integrated into the human genome

**DOI:** 10.1101/2020.12.12.422516

**Authors:** Liguo Zhang, Alexsia Richards, Andrew Khalil, Emile Wogram, Haiting Ma, Richard A. Young, Rudolf Jaenisch

## Abstract

Prolonged SARS-CoV-2 RNA shedding and recurrence of PCR-positive tests have been widely reported in patients after recovery, yet these patients most commonly are non-infectious^1–14^. Here we investigated the possibility that SARS-CoV-2 RNAs can be reverse-transcribed and integrated into the human genome and that transcription of the integrated sequences might account for PCR-positive tests. In support of this hypothesis, we found chimeric transcripts consisting of viral fused to cellular sequences in published data sets of SARS-CoV-2 infected cultured cells and primary cells of patients, consistent with the transcription of viral sequences integrated into the genome. To experimentally corroborate the possibility of viral retro-integration, we describe evidence that SARS-CoV-2 RNAs can be reverse transcribed in human cells by reverse transcriptase (RT) from LINE-1 elements or by HIV-1 RT, and that these DNA sequences can be integrated into the cell genome and subsequently be transcribed. Human endogenous LINE-1 expression was induced upon SARS-CoV-2 infection or by cytokine exposure in cultured cells, suggesting a molecular mechanism for SARS-CoV-2 retro-integration in patients. This novel feature of SARS-CoV-2 infection may explain why patients can continue to produce viral RNA after recovery and suggests a new aspect of RNA virus replication.

## Introduction

Continuous or recurrent positive SARS-CoV-2 PCR tests have been reported in patients weeks or months after recovery from an initial infection^1–14^. Although *bona fide* re-infection of SARS-CoV-2 after recovery has been reported lately^15^, cohort-based studies with strict quarantine on subjects recovered from COVID-19 suggested “re-positive” cases were not caused by re-infection^16,17^. Furthermore, no replication-competent virus was isolated or spread from these PCR-positive patients^1–3,5,6,12^. The cause for such prolonged and recurrent viral RNA production is unknown. As positive-stranded RNA viruses, SARS-CoV-2 and other beta-coronaviruses such as SARS-CoV-1 and MERS employ an RNA-dependent RNA polymerase to replicate their genomic RNA and transcribe their sub-genomic RNAs^18–20^. One possibility is that SARS-CoV-2 RNAs could be reverse-transcribed and integrated into the human genome, and transcription of the integrated DNA copies could be responsible for positive PCR tests.

Endogenous reverse transcriptase (RT) activity has been observed in human cells, and the products of reverse transcription have been shown to become integrated into the genome^21,22^. For example, *APP* transcripts have been shown to be reverse-transcribed by endogenous RT, with resultant APP fragments integrated into the genome of neurons and transcribed^22^. Human LINE-1 elements (~17% of the human genome), a type of autonomous retrotransposons, are a potential source of endogenous RT, able to retro-transpose themselves and other non-autonomous elements such as Alu^21,23^.

## Results

### Expression of viral-cellular chimeric transcripts in infected cultured and in patient-derived cells is consistent with genomic integration of viral sequences

To investigate the possibility of viral integration into virus infected cells we analyzed published RNA-Seq data from SARS-CoV-2-infected cells for evidence of chimeric transcripts, which would be indicative of viral integration into the genome and expression. Examination of these data sets ^24–30^ (Fig. S1a-b) revealed a substantial number of host-viral chimeric reads (Fig. 1a-c, S1c). These occurred in multiple sample types, including cells and organoids from lung/heart/brain/stomach tissues, as well as BALF cells directly isolated from COVID-19 patients (Fig. 1c). Chimeric read abundance was positively correlated with viral RNA level across the sample types (Fig. 1c). Chimeric reads generally accounted for 0.004% - 0.14% of total SARS-CoV-2 reads across the samples, with a 69.24% maximal number of reads in bronchoalveolar lavage fluid cells derived from severe COVID19 patients and near no chimeric reads from patient blood buffy coat cells (corresponding to almost no total SARS-CoV-2 reads). A majority of chimeric junctions mapped to SARS-CoV-2 nucleocapsid (N) sequence (Fig. 1d-e). This is consistent with the finding that nucleocapsid (N) RNA is the most abundant SARS-CoV-2 sub-genomic RNA^31^, and thus is most likely to be a target for reverse transcription and integration. These analyses support the hypothesis that SARS-CoV-2 RNA may retro-integrate into the genome of infected cells resulting in the production of chimeric viral-cellular transcripts.

**Figure 1.**
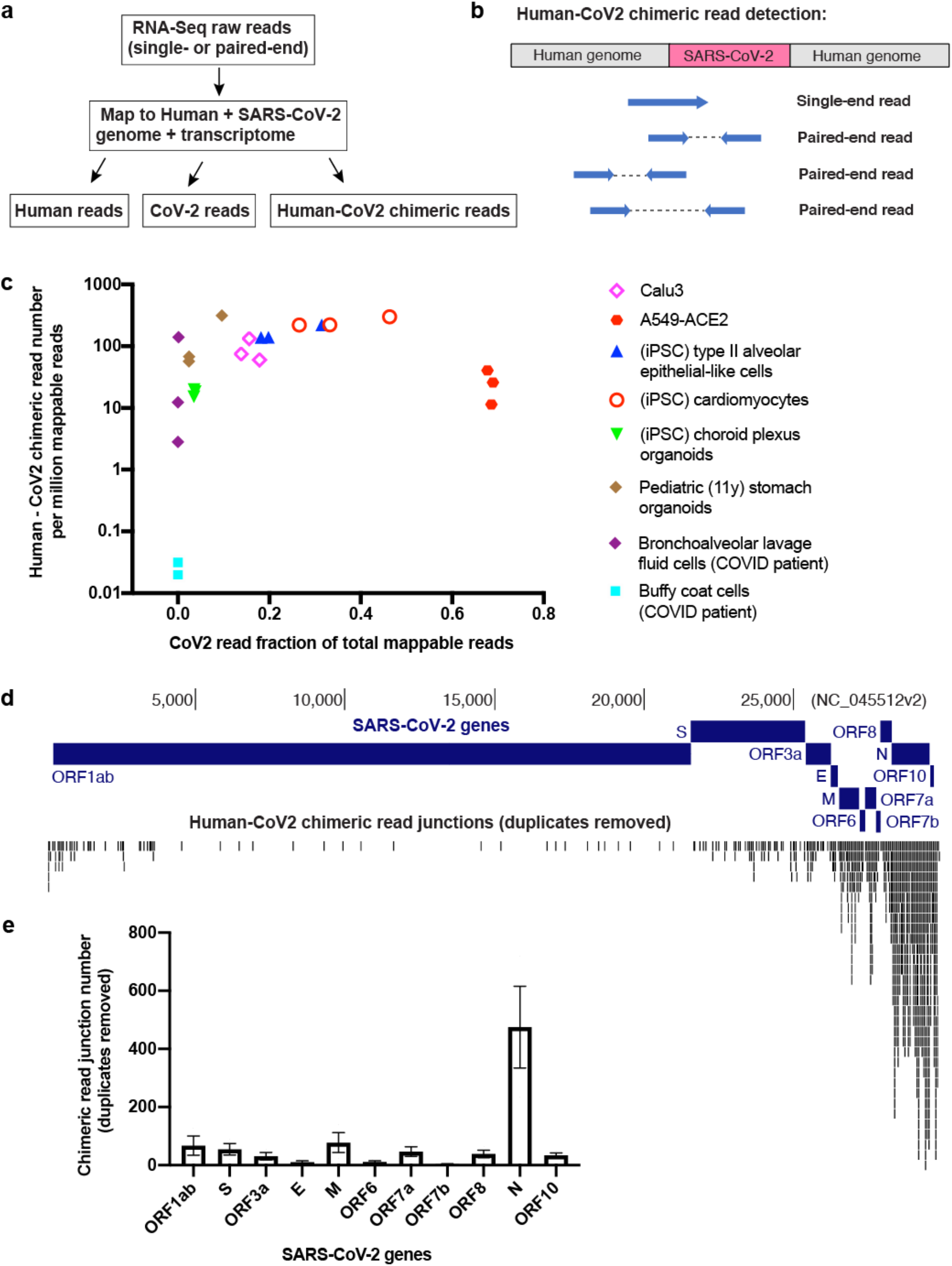
Human – SARS-CoV-2 chimeric transcripts identified in published data sets of infected cultured and patient-derived cells. **a)** Pipeline to identify human-CoV2 chimeric RNA-Seq reads. **b)** Diagram showing human-CoV2 chimeric reads mapped to potential SARS-CoV-2 integration sites in the human genome from published RNA-Seq data. **c)** Scatter plot showing human-CoV2 chimeric read number (per million total mappable reads, y-axis) versus SARS-CoV-2 read fraction of total mappable reads (x-axis) in published RNA-Seq datasets (summarized in **Supplementary Figure 1a**) from different bio-samples with SARS-CoV-2 infection. **d-e)** Human-CoV2 chimeric read junctions (duplicates removed) mapped to the SARS-CoV-2 genome (**d**) and distribution among SARS-CoV-2 genes (**e**, three biological replicates; mean ± s.e.m.). RNA-Seq data is from SARS-CoV-2 infected Calu3 cells (see **Supplementary Figure 1a**). Chimeric read junction is defined as the “breaking point” of sequences mapped to human or SARS-CoV-2 genome/transcriptome in a given RNA-Seq read.

### SARS-CoV-2 RNA can be reverse-transcribed and integrated into the human genome in cells overexpressing a reverse transcriptase

To provide experimental evidence for reverse-transcription and integration of SARS-CoV-2 RNA, we overexpressed human LINE-1 or HIV-1 reverse transcriptase (RT) in HEK293T cells and infected the transduced cells with SARS-CoV-2. The cells were tested 2 days after infection for viral sequences by PCR or fluorescence *in situ* hybridization (FISH) (Fig. 2a). Considering that the N RNA is the most abundant SARS-CoV-2 sub-genomic RNA^31^ and is most likely to be retro-integrated (Fig. 1d-e), we chose four N – targeting PCR primer sets that are used in COVID-19 tests (primer source from WHO^32^, Fig. 2a). PCR amplification of purified cellular DNA showed positive gel-bands in cells with human LINE-1 or HIV-1 RT overexpression (Fig. 2b) but not in non-transfected or non-infected cells. To test whether the DNA copies of N sequences were integrated into the cellular genome, we gel-purified cell genomic DNA (gDNA, >23 kb, Fig. S2a) and qPCR confirmed N sequences in gDNA of cells with expression of all three types of RT (Fig. 2c). Cells with strong expression of LINE-1 driven by a CMV promoter showed ~8-fold higher signals of N sequence detection suggesting a higher copy-number of integrated N sequences than in cells expressing LINE-1 driven by its natural promoter (5’UTR) or HIV-1 RT (Fig. 2c). We were able to clone full-length N DNA from gDNA of cells overexpressing CMV-LINE-1 and confirmed its sequence by Sanger sequencing (Fig. S2b). We did not detect the full-length N sequence from gDNA of cells transfected with 5’UTR-LINE-1 or HIV-1 RT, which may be due to lower expression of RT in these cells (Fig. S2b). We further confirmed that purified SARS-CoV-2 RNA from infected cells can be reverse-transcribed *in vitro* by lysates of cells expressing either LINE-1 or HIV-1 RT (Fig. S2c-d).

**Figure 2.**
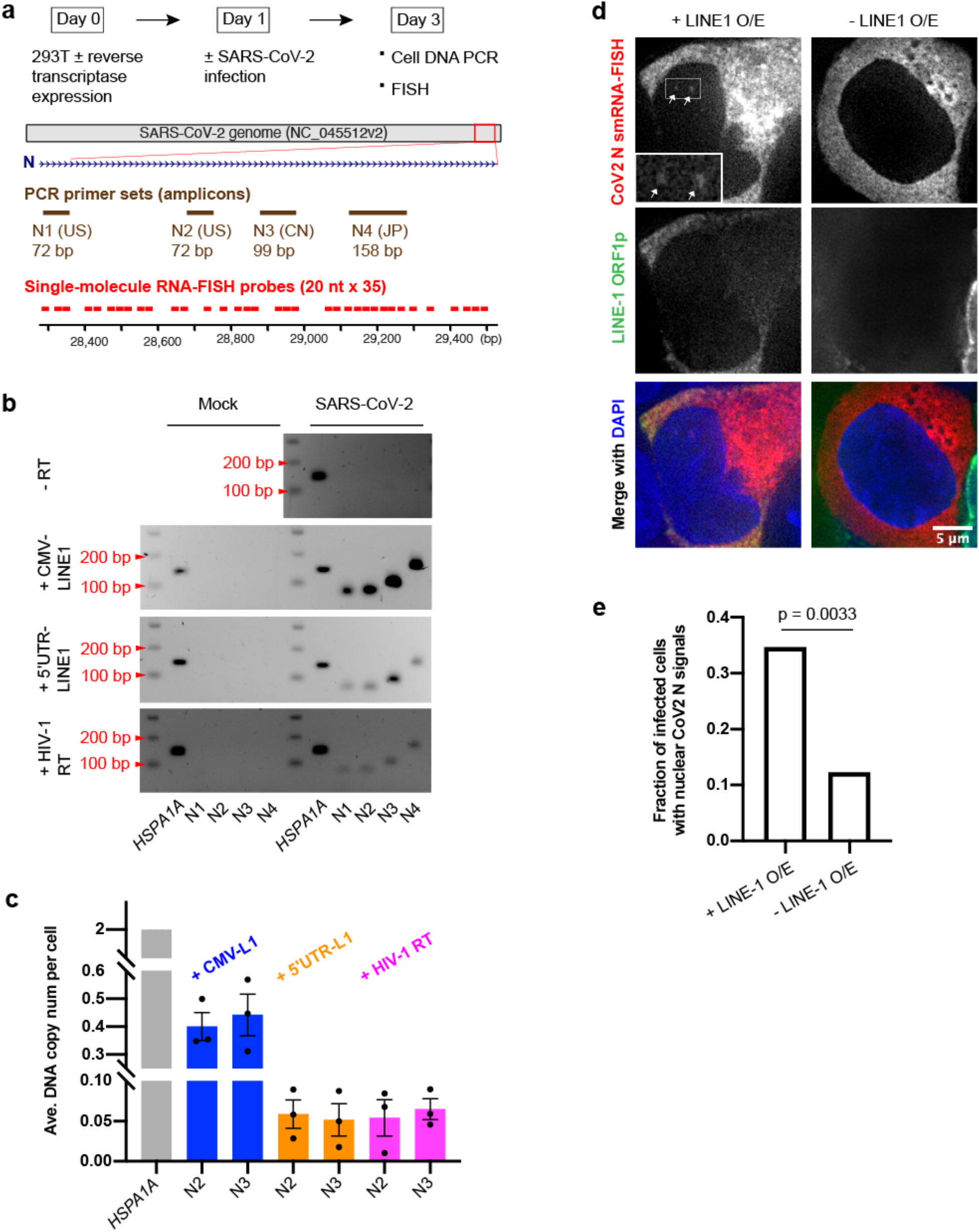
SARS-CoV-2 RNA can be reverse-transcribed and integrated into the host genome in cells with reverse transcriptase expression. **a)** Experimental workflow (top), PCR primer sets (shown as amplicons, brown) and single-molecule RNA-FISH probes (red) to detect reverse-transcription and integration of SARS-CoV-2 nucleocapsid (N) sequence (middle, blue). **b)** PCR detection of SARS-CoV-2 N sequences in cellular DNA purified from mock (left) or SARS-CoV-2 (right) infected HEK293T cells without or with transfection of human LINE-1 (CMV-LINE1 or 5’UTR-LINE1) or HIV-1 RT. *HSPA1A*: human *HSPA1A* gene as control; N1 – N4: SARS-CoV-2 N sequences as shown in **a)**. **c)** qPCR detection and copy-number estimation of SARS-CoV-2 N sequences on gel-purified HEK293T genomic DNA. *HSPA1A*: human *HSPA1A* gene as a reference; N2, N3: SARS-CoV-2 N sequences as shown in **a)**. Three biological replicates; mean ± standard error of the mean (s.e.m.). **d)** Single-molecule RNA-FISH (red) targeting SARS-CoV-2 N sequence using probes shown in **a)** plus LINE-1 ORF1 protein immuno-staining (green) and merged channels with DAPI (blue) in SARS-CoV-2 infected HEK293T cells with (left) or without (right) transfected LINE-1. Insets: 2.5x enlargement of region in white-box to show nuclear signals of SARS-CoV-2 N sequence (white arrows). Images were single z-slices from 3D optical sections (0.2-μm z-steps). **e)** Fraction of HEK293T cells infected by SARS-CoV-2 (indicated by cytoplasmic FISH signals) showing nuclear FISH signals of N sequence with (+ LINE-1 O/E, n = 75) or without (-LINE-1 O/E, n = 57) LINE-1 overexpression (indicated by LINE-1 ORF1 protein immuno-staining). Combination of two independent cell samples; Chi-Square Test of Homogeneity.

We conducted single-molecule RNA-FISH (smRNA-FISH) using fluorophore-labeled oligo-nucleotide probes targeting N (Fig. 2a) to confirm that viral N sequences were integrated and detected their transcription in the nucleus. SARS-CoV-2 infected cells showed the expected cytoplasmic FISH signals of N RNA (Fig. S3a). N RNA FISH signals were detected in cell nuclei with cells overexpressing LINE-1 (Fig. 2d, S3b), indicating nascent transcription sites of integrated N sequences. In the same cell population, a significantly higher fraction (~35%) of infected cells overexpressing LINE-1, as indicated by LINE-1 ORF1p immunostaining, showed nuclear N signals than cells not overexpressing LINE-1 (~12%) (Fig. 2e). A significantly higher fraction of infected cells that were transfected with LINE-1 plasmid (~80% transfection efficiency) showed positive nuclear N FISH signals (~30%) as compared to non-transfected cells (13%; Fig. S3c). Infected but not transfected cells also exhibited nuclear N signals, albeit at a lower frequency (~10%; Fig. 2e, S3c), implying integration of SARS-CoV-2 N RNA by cell endogenous RT activity.

### Human endogenous LINE-1 expression induced by SARS-CoV-2 infection and cytokines correlates with retro-integration

Human LINE-1 elements are autonomous retro-transposons with their encoded reverse transcriptase (ORF2p) and supporting protein (ORF1p) also aiding non-autonomous elements to retro-transpose, such as Alu and other cellular RNAs^21^. We found that expression of LINE-1 elements was significantly up-regulated in published RNA-Seq data of cells upon infection with SARS-CoV-2 and correlated with chimeric read abundance (Fig. 3a-b, S4a-d, compare Calu3 cells that are efficiently infected versus NHBE cells that are resistant to infection). Although the upregulation in Calu3 was not higher than that in NHBE, multiple LINE-1 elements were upregulated as compared to just one in NHBE (Fig. 3a, S4b, d). Expression analysis using LINE-1 specific primers^33,34^ showed a ~3-4-fold up-regulation of LINE-1 in Calu3 cells when infected by SARS-CoV-2 (Fig. 3c). Moreover, PCR analysis on Calu3 cellular DNA showed retro-integration of SARS-CoV-2 N sequences after infection (Fig. 3d-e), possibly by the activated LINE-1 reverse transcriptase.

**Figure 3.**
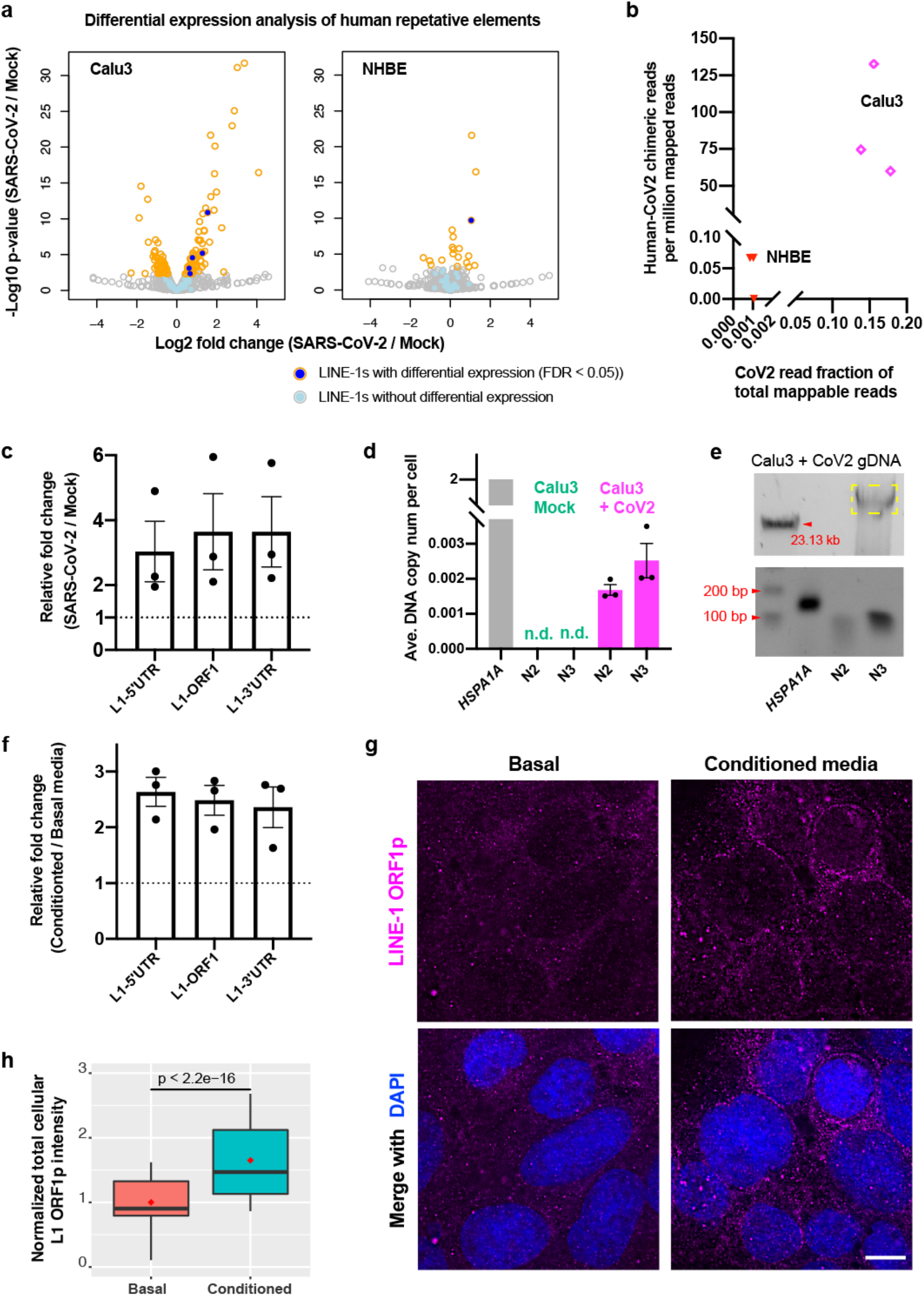
LINE-1 expression as an endogenous reverse-transcriptase source in human cells is induced by SARS-CoV-2 infection and cytokine-containing conditioned media treatment. **a)** RNA-Seq (GSE147507, see **Supplementary Figure 1a**) differential expression analysis for all human repetitive elements in SARS-CoV-2 versus mock-infected Calu3 (left) or NHBE (right) cells. Volcano plots showing −Log10 p-values (y-axis) versus Log2 fold-changes (x-axis) for all human repetitive elements with (orange circle) or without (grey circle) significant expression changes (SARS-CoV-2 versus mock-infected); dots: LINE-1 families with (dark blue) or without (light blue) significant expression changes. **b)** Scatter plot showing human-CoV2 chimeric read number (per million total mappable reads, y-axis) versus SARS-CoV-2 read fraction of total mappable reads (x-axis) in published RNA-Seq (GSE147507, see **Supplementary Figure 1a**) from infected Calu3 (magenta) or NHBE (red) cells. **c)** Endogenous LINE-1 expression fold-changes between SARS-CoV-2 versus mock-infected Calu3 cells measured by RT-qPCR with primers probing 5’UTR, ORF1, or 3’UTR regions of LINE-1. Reference genes: *GAPDH* and *TUBB*. Three biological replicates; mean ± s.e.m. **d)** qPCR detection and copy-number estimation of SARS-CoV-2 N sequences in mock (green) or SARS-CoV-2 infected (magenta) Calu3 cellular DNA. *HSPA1A*: human *HSPA1A* gene as a reference; N2, N3: SARS-CoV-2 N sequences as shown in **Figure 1a**. Three biological replicates; mean ± s.e.m; n.d.: not detected. **e)** Gel purification of large-fragment genomic DNA (yellow box, top) from SARS-CoV-2 infected Calu3 cells and PCR detection of SARS-CoV-2 N sequences in the purified genomic DNA (bottom) with same primer sets as in **d)**. **f)** Endogenous LINE-1 expression fold-changes in Calu3 cells comparing Myeloid conditioned versus basal media treatment measured by RT-qPCR with primers probing 5’UTR, ORF1, or 3’UTR regions of LINE-1. Reference genes: *GAPDH* and *TUBB*. Three biological replicates; mean ± s.e.m. **g)** LINE-1 ORF1 protein immuno-staining (magenta, same exposure and intensity scaling) plus merged channels with DAPI (blue) in Calu3 cells cultured in basal or myeloid conditioned media. Scale bar: 10 μm. **h)** Normalized cellular total LINE-1 ORF1p immuno-staining signals of Calu3 cells cultured in basal (n = 84, mean = 1.0, median = 0.9) or myeloid conditioned media (n = 126, mean = 1.7, median = 1.5). Combination of two independent cell samples. Box plots show median (inside line), means (red dot), interquartile range (IQR, box), and upper/lower quartile ± 1.5-times IQRs (whiskers). Welch’s t-test.

Patients infected with SARS-CoV-2 and other corona viruses show evidence of cytokine induction associated with the immune response, and in severe cases experience a cytokine storm^35–37^, prompting us to investigate whether cytokines alone can induce LINE-1 activation. We treated cells with cytokine-containing conditioned media from Myeloid, Microglia, or CAR-T cell cultures and found a ~2-3-fold upregulation of endogenous LINE-1 expression by PCR analysis (Fig. 3f, S5b). Expressed LINE-1 protein (ORF1p) was also confirmed by immunofluorescence staining (Fig. 3g-h, S5a). In summary, our results show induced LINE-1 expression in cells stressed by viral infection or exposed to cytokines, suggesting a molecular mechanism for SARS-CoV-2 retro-integration in human cells.

## Discussion

In this study, we showed evidence that SARS-CoV-2 RNAs can be reverse-transcribed and integrated into the human genome by several sources of reverse transcriptase such as activated human LINE-1 or co-infected retrovirus (HIV). We found LINE-1 expression can be induced upon SARS-CoV-2 infection or cytokine exposure, suggesting a molecular mechanism responsible for SARS-CoV-2 retro-integration in patients. Moreover, our results suggest that the integrated SARS-CoV-2 sequences can be transcribed, as shown by RNA-Seq and smRNA-FISH data, providing a possible explanation for the presence of viral sequences at later times after initial virus exposure and in the absence of detectable infectious virus^1–14^. The retro-inserted SRAS-CoV-2 sequences are most likely sub-genomic fragments, as the integration junctions are mostly enriched at the N sequence (Fig. 1d-e), excluding the production of infectious virus. Our data may also explain that patients, after recovery from disease symptoms, may become again positive for viral sequences as detected by PCR^1,8–14^.

An important follow-up question is whether these integrated SARS-CoV-2 sequences can express viral antigens. If so, it will be of clinical interest to assess whether viral antigens expressed from integrated virus fragments could trigger an immune response in patients that could affect the course and treatment of the disease. It is possible that the clinical consequences of the integrated viral fragments may depend on their insertion sites in the human genome, and on epigenetic regulation which has been shown in HIV patients^38^. Careful analysis on SARS-CoV-2 retro-integration sites in patient samples and correlation with disease severity will help to elucidate potential clinical consequences. Furthermore, immune response may vary depending on an individual’s underlying conditions. More generally, our results suggest a novel aspect of infection possibly also for other common disease-causing RNA viruses such as Dengue, Zika or Influenza virus, which could be subject to retro-integration and perhaps affect disease progression.

Human LINE-1 accounts for ~17% of the human genome, ~100 out of 500,000 copies of which are active^21,23^. LINE-1 – encoded reverse-transcriptase (ORF2p) and supporting protein (ORF1p) are known to retro-transpose not only LINE-1 transcripts (in *Cis*), but also other RNA species such as Alu (SINE) and cellular mRNA (in *Trans*, creating processed pseudogenes), with a “target-site – primed reverse transcription” mechanism^21^. LINE-1 proteins have been shown as nucleic acid chaperones with high RNA binding affinity^39^, therefore it is perhaps not surprising that they can retro-integrate exogenous viral RNAs. From an evolutionarily perspective, retro-integration of viral RNA by LINE-1 could be an adaptive response by the host to provide sustaining antigen expression possibly enhancing protective immunity. Conversely, retro-integration of viral RNAs could be detrimental and cause a more severe immune response in patients such as a “cytokine storm” or auto-immune reactions.

Our results may also be relevant for current clinical trials of antiviral therapies^40^. The reliance of PCR tests to assess the effect of treatments on viral replication and viral load may not reflect the efficacy of the treatment to suppress viral replication as the PCR assay may detect viral transcripts from viral sequences stably integrated into the genome rather than infectious virus.

## Methods

### Cell culture and plasmid transfection

HEK293T cells were obtained from ATCC (CRL-3216) and cultured in DMEM supplemented with 10% heat-inactivated FBS (Hyclone, SH30396.03) and 2mM L-glutamine (MP Biomedicals, IC10180683) following ATCC’s method. Calu3 cells were obtained from ATCC (HTB-55) and cultured in EMEM (ATCC 30-2003) supplemented with 10% heat-inactivated FBS (Hyclone, SH30396.03) following ATCC’s method.

Plasmid for HIV-1 reverse transcriptase expression: pCMV-dR8.2 dvpr was a gift from Bob Weinberg (Addgene plasmid # 8455; http://n2t.net/addgene:8455; RRID:Addgene_8455)^41^. Plasmids for human LINE-1 expression: pBS-L1PA1-CH-mneo (CMV-LINE-1) was a gift from Astrid Roy-Engel (Addgene plasmid # 51288; http://n2t.net/addgene:51288; RRID:Addgene_51288)^42^; EF06R (5’UTR-LINE-1) was a gift from Eline Luning Prak (Addgene plasmid # 42940; http://n2t.net/addgene:42940; RRID:Addgene_42940)^43^. Transfection was done with Lipofectamine™ 3000 (Invitrogen L3000001) following manufacturer’s protocol.

### SARS-CoV-2 infection

SARS-CoV-2 USA-WA1/2020 (Gen Bank: MN985325.1) was obtained from BEI Resources and expanded and tittered on Vero cells. Cells were infected in DMEM +2% FBS for 48 hrs using multiplicity (MOI) of 0.5 for infection of HEK293T cells and an MOI of 2 for Calu3 cells. All sample processing and harvest with infectious virus were done in the BSL3 facility at the Ragon Institute.

### Nucleic acids extraction, *in vitro* reverse transcription and PCR/qPCR

DNA extraction was following a published protocol^22^. For purification of genomic DNA, extracted total cellular DNA was run on 0.4% (w/v) agarose/1x TAE gel for 1.5 hrs with a 3V/cm voltage, with λ DNA-HindIII Digest (NEB N3012S) as size markers. Large fragment bands (>23.13 kb) were cut off, frozen in −80 °C and then crushed by a pipette tip. 3 times of volume (v/w) of high T-E buffer (10 mM Tris – 10 mM EDTA, pH 8.0) was added and then NaCl was added to 200 mM. Gel solution was heated at 70 °C for 15 mins with constant mixing and then extracted with Phenol:Chloroform:Isoamyl Alcohol (25:24:1, v/v) (Life Technologies 15593031) and Chloroform:Isoamyl alcohol 24:1 (Sigma C0549-1PT). DNA was then precipitated by sodium acetate and isopropyl alcohol. For small amount of DNA, glycogen (Life Technologies 10814010) was added as a carrier to aid precipitation.

RNA extraction was done with either TRIzol™ LS Reagent (Invitrogen 10296010) or RNeasy Plus Micro Kit (Qiagen 74034) following manufacturers’ protocols. RNA reverse transcription was done with either SuperScript™ III First-Strand Synthesis SuperMix (oligo dT + random hexamer, Invitrogen 18080400) or qScript cDNA SuperMix (QuantaBio 95048-500), following manufacturers’ protocols. *In vitro* reverse transcription assay for viral RNA by cell lysates was done following a published protocl^22^.

PCR was done using AccuPrime Taq DNA Polymerase, high fidelity (Life Technologies 12346094). qPCR was done using SYBR™ Green PCR Master Mix (Applied Biosystems 4309155) or PowerUp™ SYBR™ Green Master Mix (Applied Biosystems A25742) in a QuantStudio™ 6 system (Applied Biosystems). See **Supplementary Table 1** for primer sequences used in this study. qPCR plots were generated with Prism 8 (Prism).

### Immuno-fluorescence staining and single-molecule RNA-FISH

Cells subject to SARS-CoV-2 infection were grown in μ-Slide 8 Well (#1.5 polymer, Ibidi 80826) and fixed with 4% paraformaldehyde/CMF-PBS at room temperature (RT) for 30 mins. Otherwise, cells were grown on 12 mm round coverslips (#1.5, Warner Instruments 64-0712) and fixed with 1.6% paraformaldehyde/CMF-PBS at room temperature (RT) for 15 mins. Cells were permeabilized with 0.5% (v/v) Triton X-100/PBS, blocked with 4% (w/v) BSA/CMF-PBS at RT for 1 hr, incubated with 1:200 diluted anti-LINE-1 ORF1p mouse monoclonal antibody (clone 4H1, Sigma MABC1152, Lot 3493991), and then with 1:400 diluted Donkey-anti-Mouse-Alexa Fluor 594 second antibody (Invitrogen 21203).

Single-molecule RNA-FISH probes (Stellaris®) were ordered from LGC Biosearch Technologies with Quasar® 670 Dye labeling. See **Supplementary Table 2** for probe sequences. FISH procedure combining with immuno-fluorescence staining was following previous publications^44,45^.

Cells in μ-Slide were mounted with Ibidi Mounting Medium With DAPI (Ibidi 50011). Cells on coverslips were mounted with VECTASHIELD® HardSet™ Antifade Mounting Medium with DAPI (Vector Laboratories H-1500-10).

### Microscopy and imaging analysis

3D optical sections were acquired with 0.2-μm z-steps using a DeltaVision Elite Imaging System microscope system with a 100 × oil objective (NA 1.4) and a pco.edge 5.5 camera and DeltaVision SoftWoRx software (GE Healthcare). Image deconvolution was done using SoftWoRx. All figure panel images were prepared using FIJI software (ImageJ, NIH) and Adobe Illustrator 2020 (Adobe), showing deconvolved single z-slices.

To measure the LINE-1 ORF1p immuno-staining signal intensity, we projected cell optical sections (sum, 42 slices) with the “z projection” function in FIJI. We measured the sum of intensity of the entire cell area in the z-projected image as the signal intensity, subtracted the background intensity outside of cells and then divided by the mean of the “Basal media treatment” group to have the normalized signal intensity, as previously described^44,45^. All images from the same experiment were using the same exposure time and transmitted exciting light. All intensity measurements were done with non-deconvolved raw images. Box plot was done in R (version 4.0.3)^46^.

### RNA-Seq data analysis

RNA-Seq data were downloaded from GEO with the accession numbers GSE147507^24^, GSE153277^25^, GSE156754^26^, GSE157852^27^, GSE153684^28^, GSE145926^29^, GSE154998^30^ (summarized in **Supplementary Figure 1a**).

To identify human – SARS-CoV-2 chimeric reads, raw sequencing reads were aligned to concatenated human and SARS-CoV-2 genomes plus transcriptomes by STAR (version 2.7.1a)^47^. Human genome version hg38 with no alternative chromosomes and gene annotation version GRCh38.97 were used. SARS-CoV-2 genome version NC_045512.2 and gene annotation (http://hgdownload.soe.ucsc.edu/goldenPath/wuhCor1/bigZips/genes/) were used. The following STAR parameters^31^ were used to call chimeric reads unless otherwise specified (**Supplementary Figure 1a**): --chimOutType Junctions SeparateSAMold WithinBAM HardClip \ --chimScoreJunctionNonGTAG 0 \ --alignSJstitchMismatchNmax −1 −1 −1 −1 \ --chimSegmentMin 50 \ --chimJunctionOverhangMin 50.

To analyze human LINE-1 expression in RNA-Seq data, a published method, RepEnrich2^48^, was used to map RNA-Seq reads to human repeat annotations, using human repeat masker (hg38). Differential expression was analyzed using EdgeR package (version 3.30.3)^49,50^ in R (version 4.0.3)^46^.

### Conditioned media production and treatment

As previously described^51^, myeloid precursors were derived from human pluripotent stem cells. Briefly, human embryonic stem cells were cultured in StemFlex (ThermoFisher) feeder-free medium on Matrigel™-(Corning) coated tissue culture polystyrene. 24 hrs before single-cell harvesting via TrypLE Express (ThermoFisher), cells were treated with 10 μM ROCK Inhibitor (Y-27632) (Stem Cell Technologies) in Essential 8 (E8) medium (ThermoFisher). After harvesting, cells were centrifuged at 300 g for 3 mins in non-adherent U-bottom 96-well plates (Corning) at 10,000 cells per 150 μL/well of embryoid body (EB) medium consisting of 10 μM ROCK Inhibitor, 50 ng/mL BMP-4 (Peprotech), 20 ng/mL SCF (Peprotech), 50 ng/mL VEGF (Peprotech), and 100 U/mL Penn/Strep (ThermoFisher) in E8 base medium. EBs were cultured in the 96-well plates for 4 days with 150 μL/well of EB medium added at day 2. After 4 days, 16 EBs/well were plated in a 6-well tissue culture polystyrene plated coated with Matrigel™ in hematopoietic myeloid medium (HIM) consisting of 2mM GlutaMax (ThermoFisher), 55 μM beta-mercaptoethanol, 100 ng/mL M-CSF (Peprotech), and 25 ng/mL IL-3 (Peprotech) in X-VIVO 15 base medium (Lonza). HIM media was changed every 3-4 days for 2-3 weeks until floating CD14-positive myeloid precursors emerged. Myeloid conditioned media consisted of floating myeloid cells cultured in HIM media for 7 days at a concentration of 0.5 × 10^6^ – 1 × 10^6^ cells/mL. Cells in conditioned media were removed by centrifugation and filtration through 0.2 μM filters. Calu3 cells were cultured in the myeloid conditioned media or HIM media (basal) for two days with daily media change before harvest or fixation.

Microglia were differentiated from human induced pluripotent stem cells (hiPSCs) via embryoid bodies and primitive macrophage precursors (PMPs)^51^. In brief, hiPSCs (cultured feeder-free on matrigel in StemFlexTM (Gibco)) were dissociated with TrypLE Express (Gibco), and 10,000 cells were plated per well in 96-well ultra-low attachment plates (Corning) in 100 μL embryoid body medium (10 μM ROCK inhibitor, 50 ng/mL BMP-4, 20 ng/mL SCF, and 50 ng/mL VEGF-121 in StemFlex), before centrifugation at 300 × g for 3 mins at 4 °C. Embryoid bodies were cultured for 4 days, with adding 100 μL embryoid body medium after 2 days. 12 to 16 embryoid bodies were plated per well of tissue culture-treated 6-well plates and cultured in 3 mL hematopoetic medium (2 mM GlutaMax, 100 U/mL penicillin, 100 μg/mL streptomycin, 55 μM β-mercaptoethanol, 100 ng/mL M-CSF, 25 ng/mL IL-3, 100 U/mL penicillin, 100 μg/mL streptomycin in X-VIVO 15 (Lonza, BW04418Q). From this point on, 2 mL medium was exchanged every 4–7 days. PMPs were harvested from suspension during medium exchange and plated in microglia differentiation media over 7-14 days to produce microglia like cell monocultures (Neurobasal (Life Technologies 21103049) supplemented with Gem21 NeuroPlex without Vitamin A (GeminiBio, 400-161), 2mM GlutaMAX (Gibco), 100 ng/mL IL-34, and 10 ng/mL GM-CSF, 100 U/mL penicillin, 100 μg/mL streptomycin). For microglia stimulation, microglia differentiation media was exchanged with HEK293T media (DMEM + 10% heat-inactivated FBS + final 2mM L-Glutamine) and supplemented with 100 hg/ml lipopolysaccharide (LPS, Sigma Aldrich L4391-1MG) or PBS. After 24 hrs, the microglia conditioned media was collected, centrifugated (1000 rpm 10min) and the supernatant was directly applied to HEK293T cells. HEK293T cells received microglia conditioned media or basal HEK293T media on three constitutive days before fixation.

Human anti-CD19 CAR-T cells were generated by transduction of primary T cells purified from human peripheral blood mononuclear cells (PBMC) with CD19-CAR expressing retrovirus^52^. Anti-CD19 CAR-T cells were co-cultured with CD19-expressing beta-like cells^52^ or WIBR3 cells with a luciferase-2A-CD19 expressing cassette integrated at the AAVS1 locus in RPMI1640 medium with 10% human serum AB. Cells in the conditioned medium were removed by filtration through 0.45 μM filters. RPMI1640 medium with 10% human serum AB was used as basal media control. Calu3 cells were cultured in the CAR-T conditioned media with indicated dilutions or in the basal media for two days before harvest.

### Data Availability

The datasets generated during and/or analysed during the current study are available from the corresponding author on reasonable request.

## Acknowledgements

We thank members in the laboratories of Rudolf Jaenisch and Richard Young and other colleagues from Whitehead Institute and MIT for helpful discussions and resources. We thank Wendy Salmon from the Whitehead W.M. Keck Microscopy Facility and M. Inmaculada Barrasa from the Whitehead Bioinformatics and Research Computing for technical advice. This work was supported by grants from the NIH to RJ (1U19AI131135-01, 5R01MH104610-21) and by a generous gift from Dewpoint Therapeutics and from Jim Stone. ASK would like to acknowledge funding from the NIH (Grant: T32 EB016652). Finally, we thank Nathans Island for inspiration.

## Author contributions

Project design by R.J. and R.A.Y, execution of experiments and data analysis by L.Z., A.R, R.J and R.A.Y; E.W., A.K., and H.M. generated cells and reagents; Manuscript preparation by L.Z. and R.J. with input from all authors.

---

**Supplementary Figure 1.**
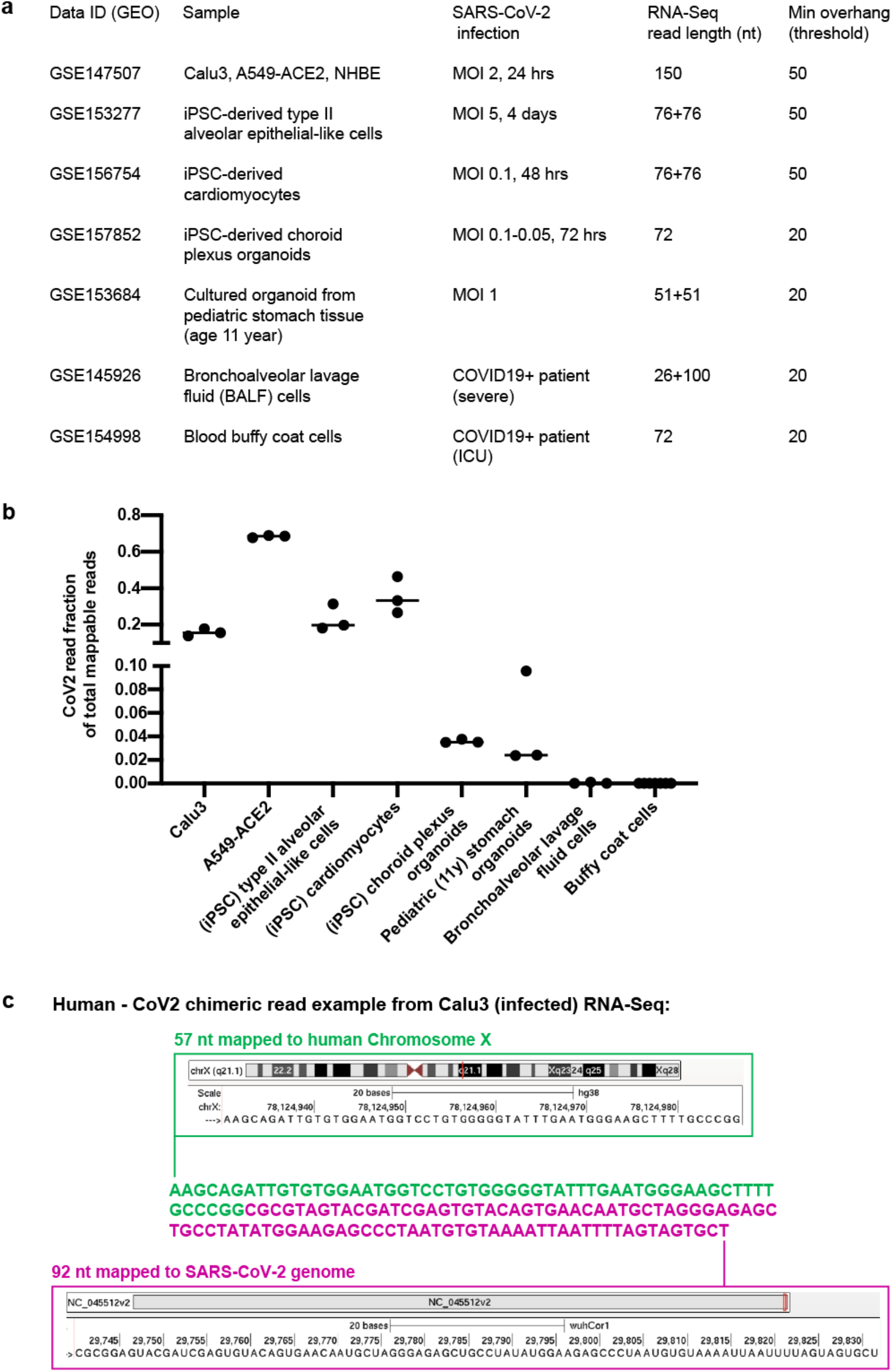
Human – SARS-CoV-2 chimeric reads identified from published RNA-Seq data. **a)** Published data used to identify human – CoV2 chimeric reads summarizing GEO accession number (data ID), sample type, infection method/type (MOI: Multiplicity Of Infection), RNA-Seq format (single or paired-end with read length), and threshold to call chimeric reads (Min overhang: minimum number of bases mapped to either human or SARS-CoV-2 genome/transcriptome to call a chimeric reads). **b)** Comparison of SARS-CoV-2 read fraction of total mappable reads in the published RNA-Seq datasets as shown in **a)**. **c)** One chimeric read example (149 nt) from Calu3 (infected) RNA-Seq with 57 nt mapped to human Chromosome × (green) and 92 nt (magenta) mapped to the SARS-CoV-2 genome.

**Supplementary Figure 2.**
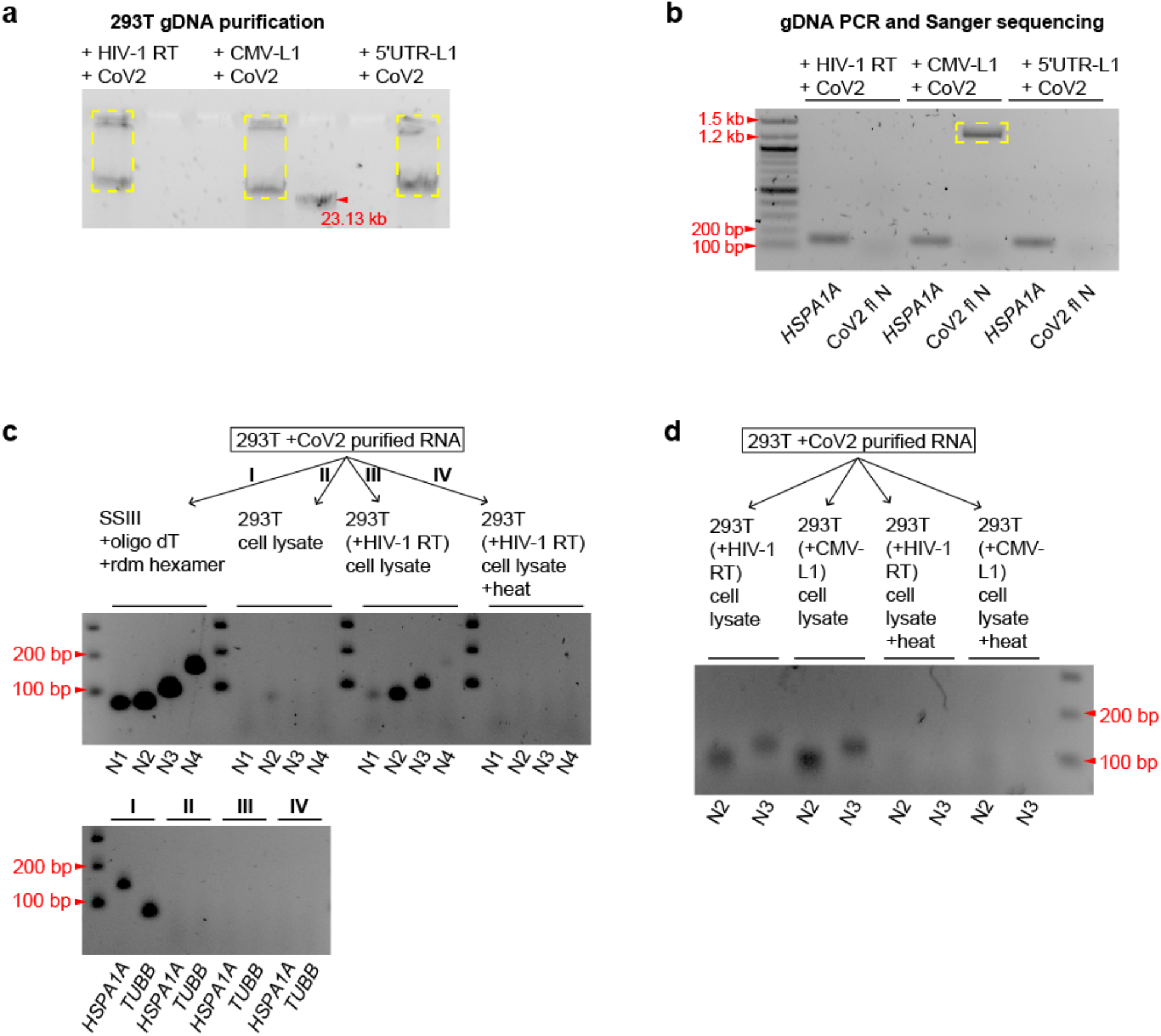
SARS-CoV-2 RNA can be reverse-transcribed *in vivo* and *in vitro* by other sources of reverse transcriptase. **a)** Gel purification of large-fragment genomic DNA (yellow boxes) from SARS-CoV-2 infected HEK293T cells carrying transfected HIV-1 RT, CMV-LINE1 or 5’UTR-LINE1. **b)** Cloning and Sanger sequencing of DNA copy of full-length SARS-CoV-2 N sequence (CoV2 fl N, yellow box) from gel-purified HEK293T genomic DNA as shown in **a)**. CoV2 fl N: amplification of full-length N sequence (1.26 kb) by primers targeting the two ends of N. *HSPA1A*: human *HSPA1A* gene as a control. Note that we can only detect full-length N sequence in gDNA from cells with CMV-LINE-1 expression, corresponding to the high copy-number of integrated N sequences as shown in **Figure 2c**. **c)***In vitro* reverse transcription of SARS-CoV-2 RNA by adding RNA purified from SARS-Cov-2 infected HEK293T cells to a commercial reverse transcriptase (I, SSIII, with oligo dT and random hexamer primers, positive control), or HEK293T cell lysate (II), or lysates of HEK293T cells expressing HIV-1 reverse transcriptase without (III) or with (IV) heat inactivation. Gel images showing PCR detection of SARS-CoV-2 N sequences from the *in vitro* reverse transcription products using primer sets (N1 – N4) as shown in **Figure 2a**. *HSPA1A* and *TUBB*: PCR primer sets against human *HSPA1A* and *TUBB* genes as controls. **d)** Same *in vitro* reverse transcription and PCR detection setup as in **c)** using lysates of HEK293T cells expressing HIV-1 reverse transcriptase or human LINE-1.

**Supplementary Figure 3.**
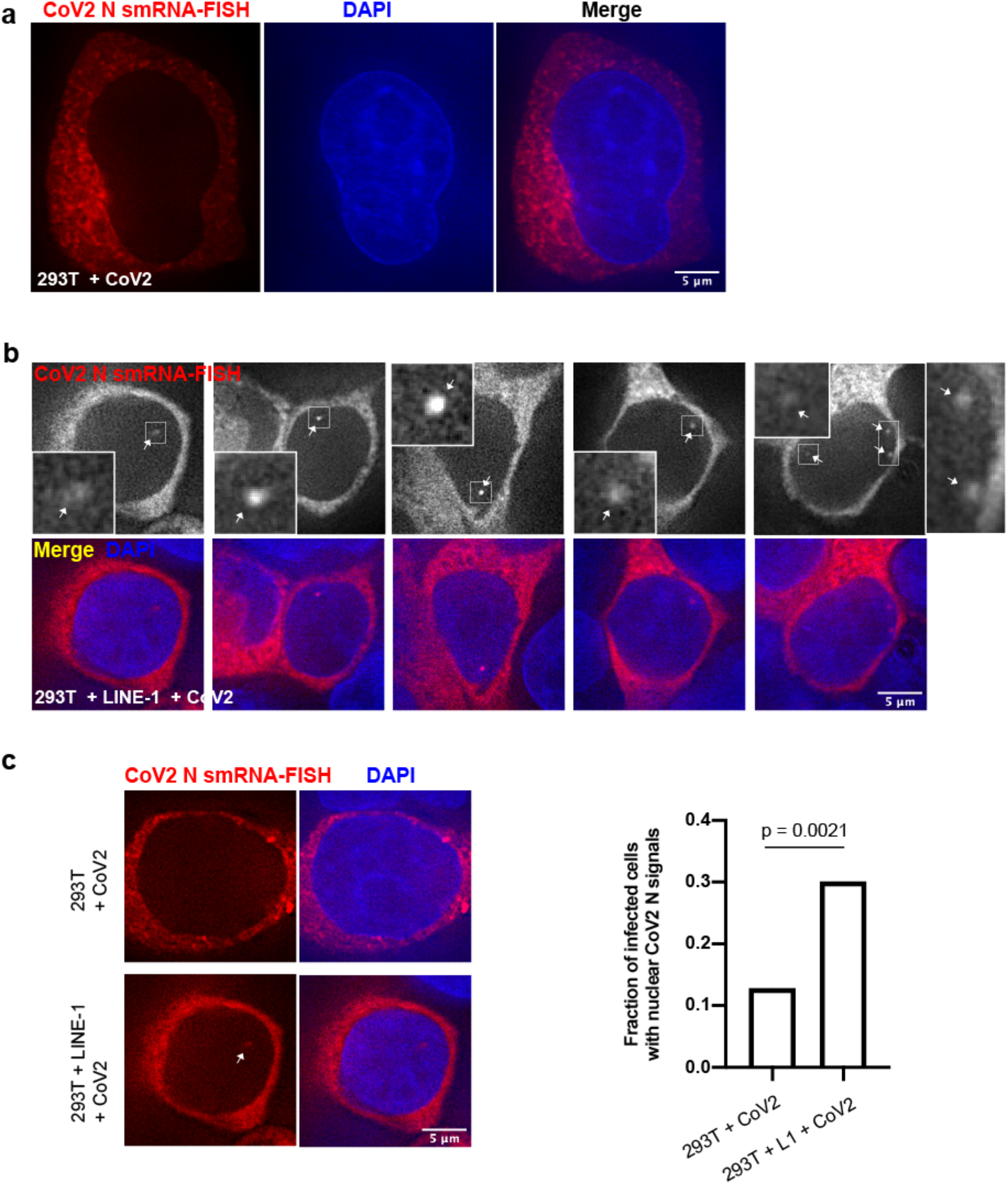
SARS-CoV-2 N RNA signals detected in cell nuclei by single-molecule RNA-FISH. **a-b)** Example images of single-molecule RNA-FISH (red/grey) targeting SARS-CoV-2 N sequence using probes shown in **Figure 2a** and merged channels with DAPI (blue) in SARS-CoV-2 infected HEK293T cells without (**a**) or with (**b**) human LINE-1 transfection. Insets in **b)**: 4x enlargement of regions in white-boxes to show nuclear signals of SARS-CoV-2 N sequence (white arrows). **c)** Comparison of nuclear N RNA-FISH signals in SARS-CoV-2 infected HEK293T cells without or with human LINE-1 transfection. Left: example images as in **a)** and **b);** Right: fraction of HEK293T cells infected by SARS-CoV-2 (indicated by cytoplasmic FISH signals) showing nuclear N RNA-FISH signals in cell populations without (left bar, n = 109) or with (right bar, n = 103) CMV-LINE-1 plasmid transfection (~80% transfection efficiency). Combination of two independent cell samples; Chi-Square Test of Homogeneity. All images shown were single z-slices from 3D optical sections (0.2-μm z-steps).

**Supplementary Figure 4.**
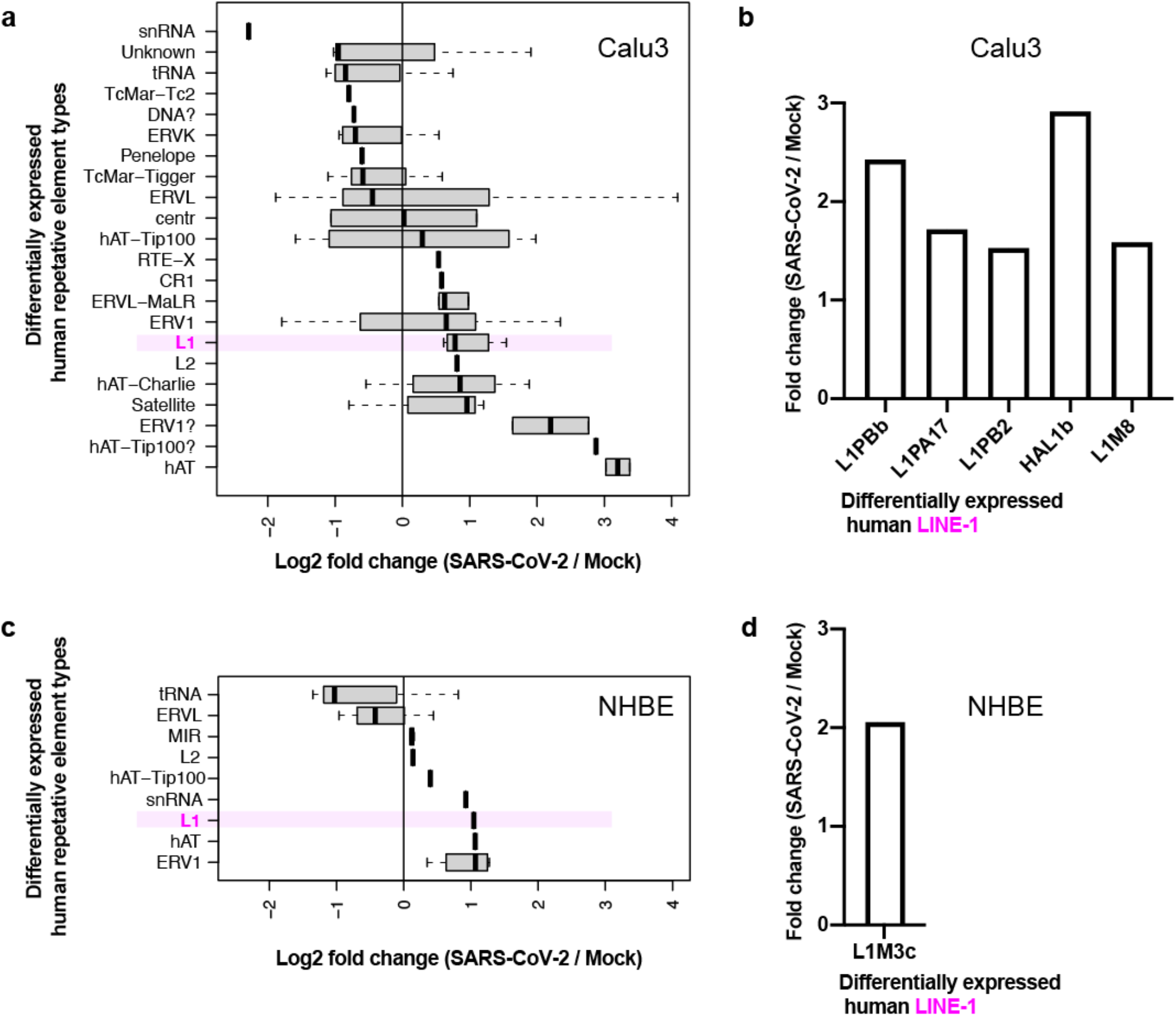
LINE-1 induction in human cells correlates with SARS-CoV-2 infection. **a, c)** Log2 fold-changes (x-axis) of different types of human repetitive elements (y-axis) with significant (FDR < 0.05) expression changes in SARS-CoV-2 versus mock infected Calu3 (**a**) or NHBE (**c**) cells from published RNA-Seq data (GSE147507). **b, d)** Fold changes (y-axis) of different human LINE-1 families (x-axis) with significant (FDR < 0.05) expression changes in SARS-CoV-2 versus mock infected Calu3 (**b**) or NHBE (**d**) cells from published RNA-Seq data (GSE147507, see **Supplementary Figure 1a**).

**Supplementary Figure 5.**
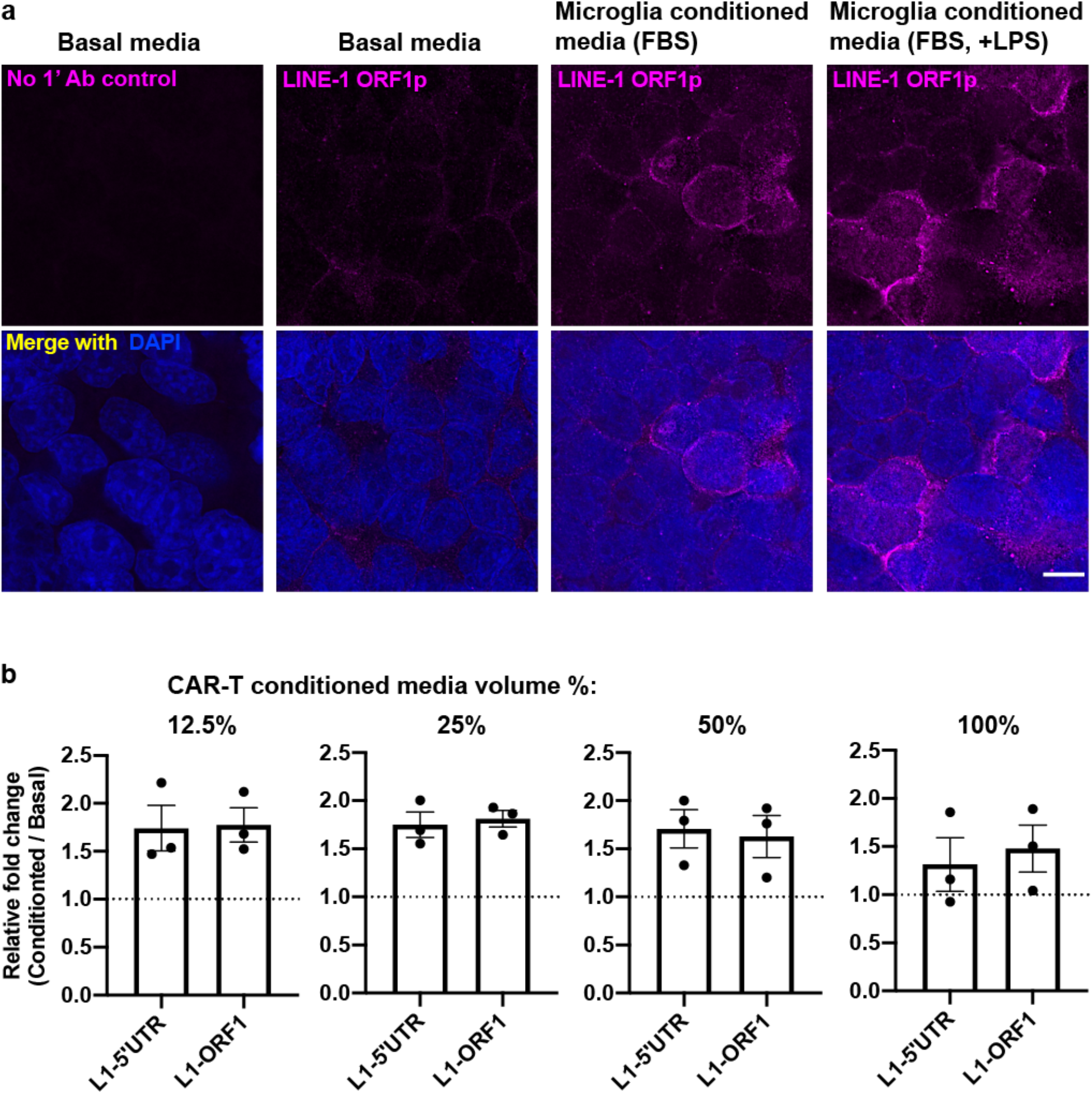
Cytokine containing media treatment triggers LINE-1 expression in human cells. **a)** LINE-1 ORF1 protein immuno-staining (magenta, same exposure and intensity scaling, 1^st^ column: no primary antibody control) plus merged channels with DAPI (blue) in HEK293T cells cultured in basal (1^st^ and 2^nd^ columns) or microglia conditioned media (3^rd^ column) or LPS-treated microglia conditioned media (4^th^ column). Scale bar: 10 μm. **b)** Endogenous LINE-1 expression fold-changes in Calu3 cells between CAR-T conditioned (diluted with basal media at indicated percentage in volume) versus basal media treatment measured by RT-qPCR with primers probing 5’UTR, ORF1, or 3’UTR regions of LINE-1. Reference genes: *GAPDH* and *TUBB*. Three independent cell samples treated with two batches of media; mean ± s.e.m.

**Supplementary Table 1.**
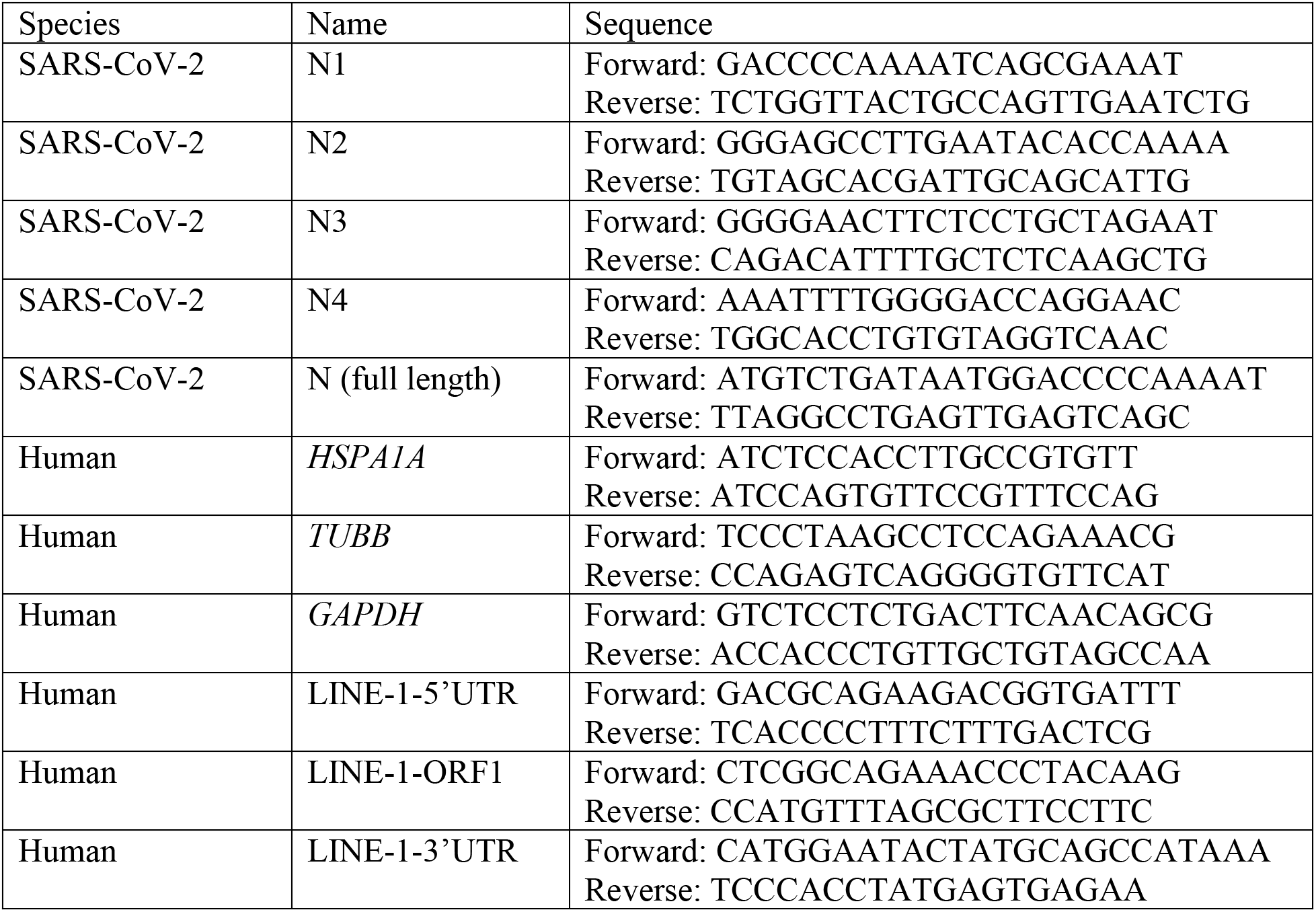
Primer sequences used in this study

**Supplementary Table 2.**
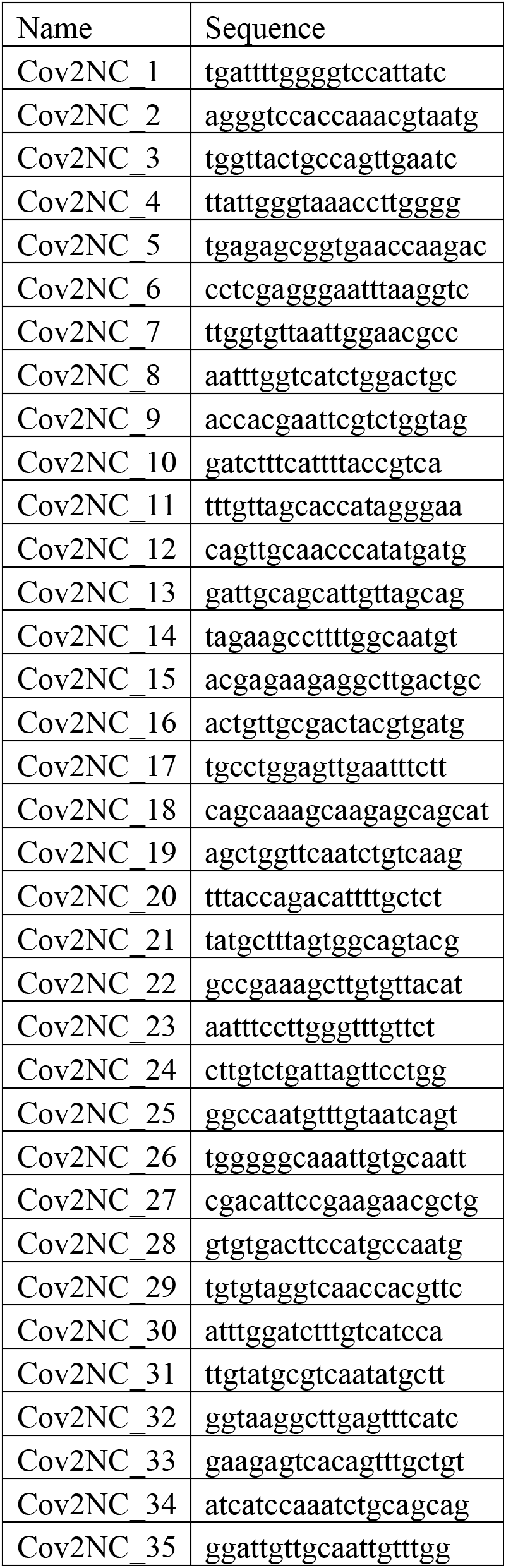
Single-molecule RNA FISH probe sequences used in this study

